# Improved neutralisation of the SARS-CoV-2 Omicron variant after Pfizer-BioNTech (BNT162b2) COVID-19 vaccine boosting with a third dose

**DOI:** 10.1101/2021.12.12.472252

**Authors:** Kerri Basile, Rebecca J Rockett, Kenneth McPhie, Michael Fennell, Jessica Johnson-Mackinnon, Jessica E Agius, Winkie Fong, Hossinur Rahman, Danny Ko, Linda Donavan, Linda Hueston, Connie Lam, Alicia Arnott, Sharon C-A Chen, Susan Maddocks, Matthew V O’Sullivan, Dominic E Dwyer, Vitali Sintchenko, Jen Kok

## Abstract

In late November 2021, the World Health Organization declared the SARS-CoV-2 lineage B.1.1.529 the fifth variant of concern, Omicron. This variant has acquired 15 mutations in the receptor binding domain of the spike protein, raising concerns that Omicron could evade naturally acquired and vaccine-derived immunity. We utilized an authentic virus, multicycle neutralisation assay to demonstrate that sera collected one, three and six months post-two doses of Pfizer-BioNTech BNT162b2 has a limited ability to neutralise SARS-CoV-2. However, four weeks after a third dose, neutralising antibody titres are boosted. Despite this increase, neutralising antibody titres are reduced four-fold for Omicron compared to lineage A.2.2 SARS-CoV-2.

## 1. Introduction

In November 2021, the SARS-CoV-2 Omicron (B.1.1.529) variant of concern (VOC) emerged in Gauteng province, South Africa, coinciding with a rapid rise in COVID-19 cases. The World Health Organization (WHO) designated Omicron a VOC on 26 November 2021 two days following its identification. [1,2] There has been subsequent worldwide spread with Omicron now the predominant circulating VOC worldwide accounting for more than 98% of sequences shared on Global Initiative on Sharing All Influenza Data (GISAID) since February 2022. [3] Early epidemiological reports from South Africa suggest that Omicron has an increased ability to evade prior infection-induced immunity compared with the Delta and Beta VOC. [1] Questions regarding Omicron’s ability to evade vaccine-derived immunity were also raised following transmission between two vaccinated individuals whilst in hotel quarantine. [2]

Vaccination plays a key role in controlling transmission of SARS-CoV-2 as well as reducing morbidity and mortality of COVID-19. The 11 COVID-19 vaccines currently granted emergency use listing (EUL) by the WHO, were all developed based on ancestral SARS-CoV-2 lineages. [4] The first to receive EUL was the Pfizer-BioNTech (BNT162b2), a nucleoside-modified RNA vaccine targeting the spike glycoprotein, the target of choice for many COVID-19 vaccines and therapeutics.

Despite the rapid worldwide roll out of vaccinations a mere twelve months following the first reported cases of COVID-19, viral evolution and the emergence of VOC threatens the success of vaccines in controlling the pandemic.

Each VOC contains a constellation of mutations in the spike glycoprotein with potential for evasion of both natural and vaccine induced immunity [5–13] and increased transmissibility. Omicron is characterised by over 30 non-synonymous mutations in the spike protein (e.g. E484A, K417N, P681H, N501Y, T478K) many within key epitopes, that provide SARS-CoV-2 an advantage over host immune responses. [12–14]

Here we present data outlining the reduced ability of sera from a COVID-19 naïve cohort collected post-two and three dose vaccination with the (Pfizer-BioNTech (BNT162b2)) vaccine to neutralise Omicron compared with the Delta and wild-type lineages; however, there was improved neutralisation after a third dose of BNT162b2 vaccine or ‘booster dose’.

## 2. Methods

*SARS-CoV-2 culture*. Upper respiratory tract specimens collected in universal transport media where SARS-CoV-2 RNA was detected by reverse transcriptase real-time polymerase chain reaction (RT-PCR) were used to inoculate VeroE6 expressing transmembrane serine protease 2 (TMPRSS2) [VeroE6/TMPRSS2; JCRB1819] cells as previously outlined.[15] TMPRSS2 expressing Vero E6 cells were used for viral isolation to prevent the emergence of advantageous mutations or deletions at or near the furin cleavage site in the spike protein which can occur when VeroE6 cells are used. [16–18]

In brief, cells were seeded at 1-3×10^4^ cells/cm^2^ whilst in the log phase of replication with Dulbecco’s minimal essential medium (DMEM) (BE12-604F, Lonza Group AG) supplemented with 9% foetal bovine serum (FBS) (10099, Gibco™) and Geneticin™ Selective Antibiotic (G418 Sulfate) 1mg/mL, (10131035, Gibco™) in Costar® 25 cm^2^ cell culture flasks (430639, Corning Inc.). The media was changed within 12 hrs for inoculation media containing 1% FBS and 1% antimicrobial agents (amphotericin B deoxycholate 25 µg/mL, penicillin 10,000 U/mL and streptomycin 10,000 µg/mL) (17-745E, Lonza Group AG) to prevent microbial overgrowth and then inoculated with 500 μL of clinical specimen into Costar® 25 cm^2^ cell culture flasks (430639, Corning Inc.). Routine testing was performed to exclude mycoplasma contamination of the cell lines and all manipulation of SARS-CoV-2 cultures were performed under biosafety level 3 conditions.

Cultures were inspected daily for cytopathic effect (CPE); the inoculum and supernatant were sampled at 96 hrs for in-house quantitative reverse transcriptase real time polymerase chain reaction (RT-qPCR) targeting the SARS-CoV-2 nucleocapsid (*N*) gene as previously described. [19,20] A decrease in the cycle threshold (Ct) from the inoculum RT-qPCR result as well as the presence of CPE was used to determine the propagation of SARS-CoV-2. Viral culture supernatant was harvested 96 hrs post-infection and clarified then stored at −80 °C in 500 μL aliquots in 2mL cryovials (72.694.406, Sarstedt, Inc.) until required. SARS-CoV-2 complete genomes were sequenced from the initial clinical specimen, positive culture supernatant, and post-neutralisation (72 hrs) to quantify genomic variations that may have developed during propagation (Supplementary Table 2).

### SARS-CoV-2 viral load quantitation by RT-qPCR

A previously described RT-PCR [19,20], targeting the *N* gene was employed to estimate the viral load of the viral inoculum and post-neutralisation viral culture. Serial (10-fold) dilutions starting at 20,000 copies/µL to 2 copies/µL of a commercially available synthetic RNA control (Wuhan-1 strain, TWIST Biosciences NCBI GenBank accession MN908947.3) were used to generate a standard curve and quantify the viral load of each culture extract. The mean Ct of biological replicates was used to calculate the viral load. A positive change in the viral load between the viral inoculum (0 hrs) and 72 hrs post-neutralisation was used to indicate viral replication.

### SARS-CoV-2 sequencing

Tiling PCR was used to amplify the entire SARS-CoV-2 genome from RNA extracts of clinical specimens using primers outlined in the Midnight sequencing protocol, the viral respiratory oligo panel (RVOP, Illumina, Inc.) (Omicron), Artic v3 primers (Delta), or a previously described long amplicon methodology (Wild-type). [19,21] Each PCR included 12.5 µL Q5 High Fidelity 2x Master Mix (New England Biolabs), 1.1 µL of either pool 1 or pool 2 10 µM primer master mix, 2.5 µL of template RNA and molecular grade water was added to generate a total volume of 25 µL. Cycling conditions were initial denaturation at 95 °C for 2 min, then 35 cycles of: 95 °C for 30 s, 65 °C for 2 min 45 s, and a final extension step of 75 °C for 10 min. Pool 1 and pool 2 amplicons were combined and purified with a 1:1 ratio of AMPureXP beads (Beckman Coulter) and eluted in 30 µL of RNAase free water. Purified products were quantified using Qubit™ 1x dsDNA HS Assay Kit (Thermo Fisher Scientific) and diluted to the desired input concentration for library preparation. Sequencing libraries were prepared using Nextera XT (Illumina, Inc.) according to the manufacturer’s respective instructions and pooled with the aim of producing 1×10^6^ reads per library. Sequencing libraries were then sequenced with paired end 76 bp chemistry on the iSeq or MiniSeq (Illumina, Inc.) platforms.

### Bioinformatic analysis

Raw sequence data were processed using an in-house quality control procedure prior to further analysis as described previously. [19,22] De-multiplexed reads were quality trimmed using Trimmomatic v0.36 (sliding window of 4, minimum read quality score of 20, leading/trailing quality of 5 and minimum length of 36 after trimming). [23] Briefly, reads were mapped to the reference SARS-CoV-2 genome (NCBI GenBank accession MN908947.3) using Burrows-Wheeler Aligner (BWA)-mem version 0.7.17, [24] with unmapped reads discarded. Average genome coverage was estimated by determining the number of missing bases (Ns) in each sequenced genome. Variant calling and the generation of consensus sequences was conducted using iVar,[25] with soft clipping over primer regions (version 1.2.1, min. read depth >10x, quality >20, min frequency threshold of 0.1). Polymorphic sites that have previously been highlighted as problematic were monitored. [26] SARS-CoV-2 lineages were inferred using Phylogenetic Assignment of Named Global Outbreak LINeages (Pango v.4.1.1 PLEARN-v1.12). [27,28]

### Post Pfizer-BioNTech (BNT162b2) vaccine sera

Sera were sourced from Australian healthcare workers caring for, or handling specimens from, individuals exposed to, or diagnosed with, SARS-CoV-2 infection enrolled in the COVID Heroes Serosurvey (http://www.covidheroes.org.au; Accessed 8 December 2021). Sera were tested upon receipt with an in-house immunofluorescence assay (IFA) against SARS-CoV-2-specific immunoglobulin A (IgA), immunoglobulin M (IgM) and immunoglobulin G (IgG) and then stored at 4 °C. [29] Fourteen sera samples were included from seven participants (Supplementary Table 1). This included samples from a vaccine-naïve individual (n=1), samples from individuals who received two doses of BNT162b2 (n=9) and individuals that received a third dose of BNT162b2 (booster dose) six months after the primary schedule (n=4). Median age was 59 years [Range 34–65] with sera collected one month, three and 6 months post-primary BNT162b2 vaccination and four weeks after the third dose of BNT162b2 (booster dose). All vaccine recipients had no documented history of prior SARS-CoV-2 infection, as confirmed by absence of SARS-CoV-2-specific nucleocapsid antibodies detected by an inhouse SARS-CoV-2 enzyme-linked immunosorbent assay on serial sampling since study enrolment. Sera was heat-inactivated at 56 °C for 30 min to inactivate complement prior to micro-neutralisation.

### Determination of 50% tissue culture infective dose (TCID_50_)

The viral 50% tissue culture infective dose (TCID_50_) was determined for each variant virus. Briefly, a passage one aliquot of virus stock was serially diluted (1×10^−2^–1×10^−7^) in virus inoculation media. Four replicates of each virus dilution were used to inoculate VeroE6/TMPRSS2 cells at 60% confluence in Costar® 24-well clear tissue culture-treated multiple well plates (3524, Corning Inc.). Plates were sealed with AeraSeal® Film (BS-25, Excel Scientific Inc.) to minimise evaporation, spillage, and well-to-well cross-contamination. Plates were inspected daily for CPE and 110 μL sampled at inoculation, 48, 72 and 96 hrs. Infections were terminated at 96 hrs based on visual inspection for CPE and used in conjunction with RT-qPCR results to determine each isolate’s TCID_50_.

### Micro-neutralisation assay

VeroE6/TMPRSS2 cells were seeded with DMEM from stocks in Costar® 96-well clear tissue culture-treated flat bottom plates (3596, Corning Inc.) at 40% confluence. Cells were incubated at 37°C with 5% CO_2_ for 12 hrs or until they reached 60% confluence. Virus stocks were diluted to 200 TCID_50_ in inoculation media. Doubling dilutions from 1:10 to 1:320 of vaccine-naïve and post BNT162b2 vaccination sera were added in equal proportions with virus in a Costar® 96 well plate (3596, Corning Inc.) and incubated for 60 min at 37 °C and 5% CO_2_ to enable virus neutralisation. After this incubation the media was removed from the cell monolayer and 100 μL of fresh media was added. Each dilution of sera was performed in duplicate per virus variant; 12 wells of uninfected cells were used per plate as a negative control. Plates were sealed with AeraSeal® Film (BS-25, Excel Scientific Inc.) to minimise evaporation, spillage, and well-to-well cross-contamination. After 60 mins of viral neutralisation a residual 110 μL was sampled from the eight naïve patient wells per virus for extraction and RT-qPCR. Plates were inspected daily for CPE with a final read recorded at 72 hrs by three scientists independently. SARS-CoV-2 in-house RT-qPCR was used to quantify the viral load post-neutralisation, with 110 μL of each dilution removed at 72 hrs to determine viral load. The 110 μL of each dilution was added to 110 μL of MagNA Pure96 External Lysis buffer (06374913001, Roche) at a 1:1 ratio in a MagNA Pure 96-well deep-well extraction plate (06241603001, Roche), covered with a MagNA Pure Sealing Foil (06241638001, Roche), and left to rest in the biosafety class two cabinet for 10 mins, a time-period shown to inactivate SARS-CoV-2 by in-house verification of a published protocol. [30] The RNA was then extracted with the MagNA Pure 96 DNA and Viral NA Small volume kit (06 543 588 001, Roche) on the MagNA Pure 96 Instrument (06541089001, Roche). Median neutralisation breakpoints (CPE) were calculated using the breakpoints for individual sera for all three SARS-CoV-2 variants.

### Statistical Analysis

Mean neutralising antibody titres (nAbT) were evaluated, and statistical significance assessed using the *t* test with a two-tailed hypothesis. Results were considered statistically significant at *p* <.05. Graphs were generated using RStudio (version 3.6.1)

## 3. Results

### Levels of neutralising antibody against different SARS-CoV-2 lineages

Neutralising antibody titres (nAbT) in sera examined four weeks after a third dose of BNT162b2 vaccine (booster dose) were higher than nAbT measured at one, three, and six months after two doses of BNT162b2 (Figure 1, Table S1). However, there was a four-fold reduction in median nAbT against Omicron in contrast to the wild-type and 1.5-fold decrease of nAbT against Delta VOC following the third dose. Median nAbT after one, three, six months following two doses of BNT162b2 vaccine for all variants were documented with titres of <10 and <20 (Figure 1, Table S1). Trends in decreasing nAbT in sera collected one, three, and six months post-two doses of BNT162b2 were observed, comparable with decreasing trends in SARS-CoV-2-specific IgG levels determined by IFA (Table S1, Figure S1). Increases in nAbT were noted four weeks after BNT162b2 boosters, with median titres of 240, 160 and 60 for wild-type, Delta and Omicron respectively. Although median nAbT responses were lower for Delta (p=0.28) and Omicron (p=0.55), this reduction did not reach statistical significance.

**Figure 1.**
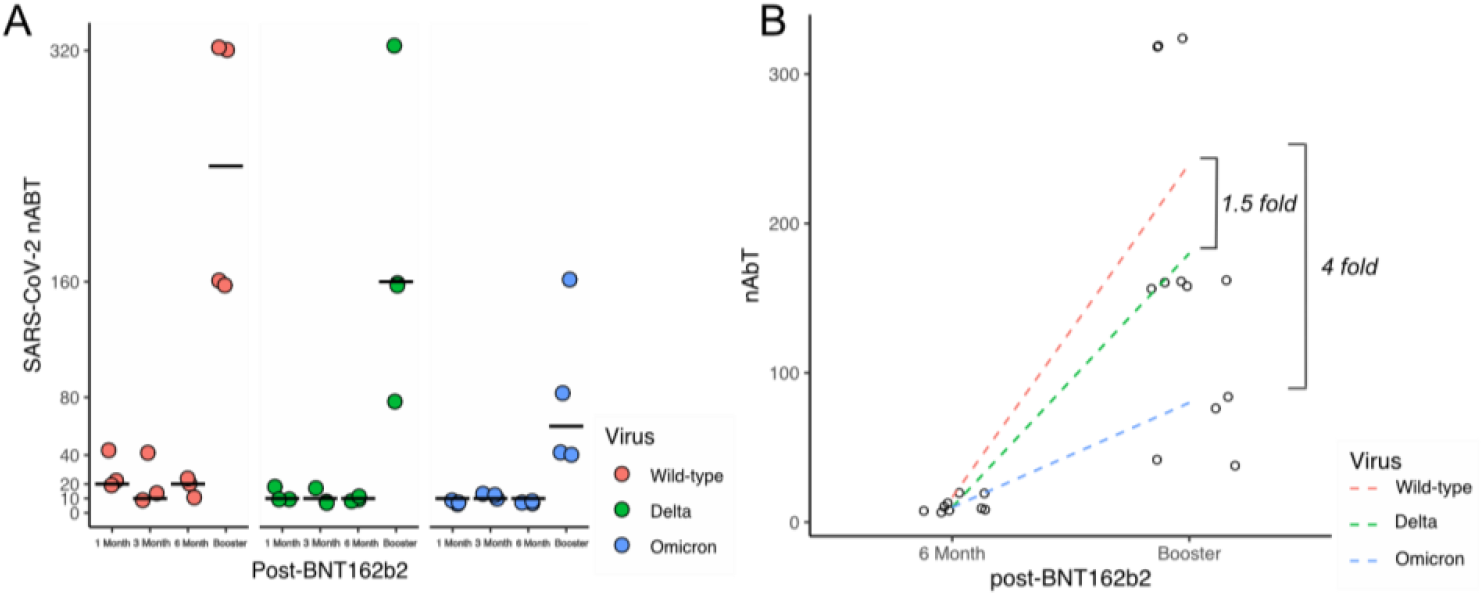
Increases in neutralising antibody titres (nAbT) four weeks after the third dose of Pfizer-BioNTech (BNT162b2) **A**. Illustrates the neutralising antibody titre (nAbT) of SARS-CoV-2 variants Delta (green) and Omicron (blue) compared to Wild-type (red, SARS-CoV-2 lineage A.2.2) black lines indicate the median titre at each timepoint. **B**. The sera from four individuals collected six months after two doses of Pfizer-BioNTech (BNT162b2) and four weeks after a third dose of BNT162b2 vaccine (booster dose) demonstrate an increase in nAbT against all three variants. However, a 1.5 and 4-fold reduction in nAbT is observed for the Delta and Omicron compared to wild-type virus. Dashed lines depict the linear regression between individual titres pre- and post-booster BNT162b2 for each virus.

### Different in vitro infection kinetics between SARS-CoV-2 lineages

All three lineages demonstrated comparable TCID_50_ and viral load at inoculation (Figure S2). Nevertheless, Omicron had a slower propagation rate with increases in viral load not detected in culture supernatant until 96 hrs post-infection (Figure S2). In contrast, wild-type and Delta cultures showed a 4-5 log_10_ increase in viral load 72 hrs post-inoculation (Figure S3).

## 4. Discussion

Omicron has rapidly evolved to encode over 30 protein substitutions in the SARS-CoV-2 spike protein, raising grave concerns for the effectiveness of COVID-19 vaccines which were designed using ancestral viruses. Although reductions in nAbT are noted when human sera from double-dose BNT162b2 vaccinated individuals were challenged with Omicron, sera after a third dose (booster dose) neutralized SARS-CoV-2. The observation of delays in Omicron growth *in vitro* suggests that these mutational changes may negatively affect viral fitness. Further investigations are required to determine if the cell infection dynamics are due to reduced efficiency in viral entry or host cell egress, although others have noted that ACE2 is still required for efficient Omicron propagation in cell culture. [31] Given the longer growth times, understanding viral kinetics of Omicron infections is important to guide infection control practices as the duration of infectivity, peak viral load and shedding may be different. Our findings also support the notion that booster doses of vaccines are likely to be required to minimise SARS-CoV-2 transmission, given that BNT162b2 is highly effective in preventing severe SARS-Cov-2 infection and mortality, particularly in those with waning immunity post primary vaccination. [32] Despite the limited sample number, this study’s strength is the serial sera samples from four participants collected pre-and post-booster with BNT162b2. This report also lacked samples from persons previously infected with SARS-CoV-2, but previous studies have demonstrated comparable antibody levels to vaccine recipients after six months. [29] Laboratory variables such as cell lines, SARS-CoV-2 variants, age groups and timing of sera collection (in relation to infection and/or vaccination) used, and test methods (TCID_50_; incubation times; type of neutralisation test [e.g. focus, plaque or micro-neutralisation]) need to be considered when interpreting nAbT studies.

## 5. Conclusions

These initial findings suggest that Omicron can be neutralized by sera collected from BNT162b2 recipients, optimally after three vaccine doses. However, the ability to neutralize Omicron is reduced, suggesting that there may be high incidences of breakthrough infections or reinfections despite vaccine boosters. Further studies with larger and targeted cohorts (e.g. young, elderly or immunocompromised) are warranted, alongside investigations into Omicron’s evasiveness to T-cell responses and primary and boosted vaccine effectiveness.

## Supplementary Materials

**Table S1:** Study Participant Demographic and In-house SARS-CoV-2 Immunofluorescence (IFA) and Neutralising Antibody Titres (nAbT); **Figure S1**. Waning levels of SARS-CoV-2-specific IgG 1, 3 and 6 months following two doses of third dose of Pfizer-BioNTech (BNT162b2) which are boosted 4 weeks after the 3 BNT162b2 dose; **Figure S2**. Infection kinetics of SARS-CoV-2 lineages propagated in VeroE6/TMPRSS2 cells; **Figure S3**. Change in SARS-CoV-2 viral load 72 hours post-neutralisation with sera collected post-Pfizer-BioNTech (BNT152b2) challenged with SARS-CoV-2 VOCs Delta and Omicron compared to wildtype (lineage A.2.2) virus; **Table S2:** Non-Silent mutations in SARS-CoV-2 isolates used in Micro-neutralisation experiments.

## Author Contributions

Conceptualization, Kerri Basile, Rebecca Rockett, Kenneth McPhie, Matthew V O’Sullivan, Dominic Dwyer and Jen Kok; Data curation, Kerri Basile, Rebecca Rockett, Jessica Johnson-Mackinnon, Jessica. Agius, Winkie Fong, Connie Lam and Alicia Arnott; Formal analysis, Kerri Basile, Rebecca Rockett and Jen Kok; Investigation, Kerri Basile, Rebecca Rockett, Michael Fennell, Hossinur Rahman, Danny Ko and Linda Donavan; Methodology, Kerri Basile, Rebecca Rockett and Kenneth McPhie; Resources, Kerri Basile, Rebecca Rockett, Hossinur Rahman, Linda Donavan, Linda Hueston, Matthew V O’Sullivan, Vitali Sintchenko and Jen Kok; Supervision, Sharon Chen, Susan Maddocks, Matthew V O’Sullivan, Dominic Dwyer, Vitali Sintchenko and Jen Kok; Visualization, Kerri Basile, Rebecca Rockett, Jessica Johnson-Mackinnon, Jessica. Agius, Winkie Fong, Connie Lam and Alicia Arnott; Writing – original draft, Kerri Basile, Rebecca Rockett and Kenneth McPhie; Writing – review & editing, Kerri Basile, Rebecca Rockett, Sharon Chen, Susan Maddocks, Matthew V O’Sullivan, Dominic Dwyer, Vitali Sintchenko and Jen Kok.

All authors have read and agreed to the published version of the manuscript.

## Funding

This study was supported by the Prevention Research Support Program funded by the NSW Ministry of Health, NSW Health COVID-19 priority funding, National Health and Medical Research Council, Australia (APPRISE 1116530). Dr Basile is supported by a Jerry Koutts PhD Scholarship from the ICPMR Trust Fund.

## Institutional Review Board Statement

Ethical and governance approval for the study was granted by the Western Sydney Local Health District Human Research Ethics Committee (2020/ETH02426) and (2020/ETH00786).

## Acknowledgments

The authors acknowledge the technical assistance provided by the Sydney Informatics Hub, a Core Research Facility of the University of Sydney, the COVID Heroes (A serosurvey of healthcare workers caring for, or handling specimens from, individuals exposed to, or diagnosed with, SARS-CoV-2 infection in Australia) study participants from where the sera were sourced, the laboratories who referred samples for testing that were included in this analysis.

## Conflicts of Interest

The authors declare no conflict of interest.

## Notes

### Competing Interest Statement

The authors have declared no competing interest.

### Summary of Updates

corrections

## References

1. Network for Genomic Surveillance in South Africa (NGS-SA) SARS-CoV-2 Sequencing Update 1 December 2021. [Internet]. Network for Genomic Surveillance in South Africa (NGS-SA); 2021. Accessed on 6 December 2021, Available at: Https://Www.Nicd.Ac.Za/Wp-Content/Uploads/2021/12/Update-of-SA-Sequencing-Data-from-GISAID-1-Dec-Final.Pdf.

2. World Health Organization Statement - Classification of Omicron (B.1.1.529): SARS-CoV-2 Variant of Concern 26 November 2021. Accessed on 6 December 2021, Available at: Https://Www.Who.Int/News/Item/26-11-2021-Classification-of-Omicron-(b.1.1.529)-Sars-Cov-2-Variant-of-Concern.

3. The World Health Organization - Tracking SARS-CoV-2 Variants. Accessed 4 August 2022, Available at: Https://Www.Who.Int/Activities/Tracking-SARS-CoV-2-Variants.

4. The World Health Organization - COVID-19 Vaccines with WHO Emergency Use Listing, Accessed 4 August 2022, Available at: Https://Extranet.Who.Int/Pqweb/Vaccines/Vaccinescovid-19-Vaccine-Eul-Issued.

5. Liu, Y.; Liu, J.; Xia, H.; Zhang, X.; Fontes-Garfias, C.R.; Swanson, K.A.; Cai, H.; Sarkar, R.; Chen, W.; Cutler, M.; et al. Neutralizing Activity of BNT162b2-Elicited Serum. New England Journal of Medicine 2021, 384, 1466–1468, doi:10.1056/nejmc2102017.

6. Lopez Bernal, J.; Andrews, N.; Gower, C.; Gallagher, E.; Simmons, R.; Thelwall, S.; Stowe, J.; Tessier, E.; Groves, N.; Dabrera, G.; et al. Effectiveness of Covid-19 Vaccines against the B.1.617.2 (Delta) Variant. New England Journal of Medicine 2021, 385, 585–594, doi:10.1056/nejmoa2108891.

7. Chemaitelly, H.; Yassine, H.M.; Benslimane, F.M.; al Khatib, H.A.; Tang, P.; Hasan, M.R.; Malek, J.A.; Coyle, P.; Ayoub, H.H.; al Kanaani, Z.; et al. MRNA-1273 COVID-19 Vaccine Effectiveness against the B.1.1.7 and B.1.351 Variants and Severe COVID-19 Disease in Qatar. Nature Medicine 2021, 27, 1614–1621, doi:10.1038/s41591-021-01446-y.

8. Graham, M.S.; Sudre, C.H.; May, A.; Antonelli, M.; Murray, B.; Varsavsky, T.; Kläser, K.; Canas, L.S.; Molteni, E.; Modat, M.; et al. Changes in Symptomatology, Reinfection, and Transmissibility Associated with the SARS-CoV-2 Variant B.1.1.7: An Ecological Study. The Lancet Public Health 2021, 6, e335–e345, doi:10.1016/S2468-2667(21)00055-4.

9. Jangra, S.; Ye, C.; Rathnasinghe, R.; Stadlbauer, D.; Alshammary, H.; Amoako, A.A.; Awawda, M.H.; Beach, K.F.; Bermúdez-González, M.C.; Chernet, R.L.; et al. SARS-CoV-2 Spike E484K Mutation Reduces Antibody Neutralisation. The Lancet Microbe 2021, 2, e283–e284.

10. Xie, X.; Liu, Y.; Liu, J.; Zhang, X.; Zou, J.; Fontes-Garfias, C.R.; Xia, H.; Swanson, K.A.; Cutler, M.; Cooper, D.; et al. Neutralization of SARS-CoV-2 Spike 69/70 Deletion, E484K and N501Y Variants by BNT162b2 Vaccine-Elicited Sera. Nature Medicine 2021, 27, 620–621, doi:10.1038/s41591-021-01270-4.

11. Willett, B.J.; Grove, J.; MacLean, O.A.; Wilkie, C.; de Lorenzo, G.; Furnon, W.; Cantoni, D.; Scott, S.; Logan, N.; Ashraf, S.; et al. SARS-CoV-2 Omicron Is an Immune Escape Variant with an Altered Cell Entry Pathway. Nature Microbiology 2022, doi:10.1038/s41564-022-01143-7.

12. VanBlargan, L.A.; Errico, J.M.; Halfmann, P.J.; Zost, S.J.; Crowe, J.E.; Purcell, L.A.; Kawaoka, Y.; Corti, D.; Fremont, D.H.; Diamond, M.S. An Infectious SARS-CoV-2 B.1.1.529 Omicron Virus Escapes Neutralization by Therapeutic Monoclonal Antibodies. Nature Medicine 2022, 28, 490–495, doi:10.1038/s41591-021-01678-y.

13. Zhang, X.; Wu, S.; Wu, B.; Yang, Q.; Chen, A.; Li, Y.; Zhang, Y.; Pan, T.; Zhang, H.; He, X. SARS-CoV-2 Omicron Strain Exhibits Potent Capabilities for Immune Evasion and Viral Entrance. Signal Transduction and Targeted Therapy 2021, 6, 430, doi:10.1038/s41392-021-00852-5.

14. Chaguza, C.; Coppi, A.; Earnest, R.; Ferguson, D.; Kerantzas, N.; Warner, F.; Young, H.P.; Breban, M.I.; Billig, K.; Koch, R.T.; et al. Rapid Emergence of SARS-CoV-2 Omicron Variant Is Associated with an Infection Advantage over Delta in Vaccinated Persons. Med 2022, 3, 325-334.e4, doi:10.1016/j.medj.2022.03.010.

15. Basile, K.; McPhie, K.; Carter, I.; Alderson, S.; Rahman, H.; Donovan, L.; Kumar, S.; Tran, T.; Ko, D.; Sivaruban, T.; et al. Cell-Based Culture of SARS-CoV-2 Informs Infectivity and Safe de-Isolation Assessments during COVID-19. Clinical Infectious Diseases 2020, doi:10.1093/cid/ciaa1579.

16. Johnson, B.A.; Xie, X.; Bailey, A.L.; Kalveram, B.; Lokugamage, K.G.; Muruato, A.; Zou, J.; Zhang, X.; Juelich, T.; Smith, J.K.; et al. Loss of Furin Cleavage Site Attenuates SARS-CoV-2 Pathogenesis. Nature 2021, 591, 293–299, doi:10.1038/s41586-021-03237-4.

17. Klimstra, W.B.; Tilston-Lunel, N.L.; Nambulli, S.; Boslett, J.; McMillen, C.M.; Gilliland, T.; Dunn, M.D.; Sun, C.; Wheeler, S.E.; Wells, A.; et al. SARS-CoV-2 Growth, Furin-Cleavage-Site Adaptation and Neutralization Using Serum from Acutely Infected Hospitalized COVID-19 Patients. Journal of General Virology 2020, 101, 1156–1169, doi:10.1099/jgv.0.001481.

18. Peacock, T.P.; Goldhill, D.H.; Zhou, J.; Baillon, L.; Frise, R.; Swann, O.C.; Kugathasan, R.; Penn, R.; Brown, J.C.; Sanchez-David, R.Y.; et al. The Furin Cleavage Site in the SARS-CoV-2 Spike Protein Is Required for Transmission in Ferrets. Nature Microbiology 2021, 6, 899–909, doi:10.1038/s41564-021-00908-w.

19. Lam, C.; Gray, K.; Gall, M.; Sadsad, R.; Arnott, A.; Johnson-Mackinnon, J.; Fong, W.; Basile, K.; Kok, J.; Dwyer, D.E.; et al. SARS-CoV-2 Genome Sequencing Methods Differ in Their Abilities To Detect Variants from Low-Viral-Load Samples. Journal of Clinical Microbiology 2021, 59, doi:10.1128/JCM.01046-21.

20. Rahman, H.; Carter, I.; Basile, K.; Donovan, L.; Kumar, S.; Tran, T.; Ko, D.; Alderson, S.; Sivaruban, T.; Eden, J.-S.; et al. Interpret with Caution: An Evaluation of the Commercial AusDiagnostics versus in-House Developed Assays for the Detection of SARS-CoV-2 Virus. Journal of Clinical Virology 2020, 127, doi:10.1016/j.jcv.2020.104374.

21. Freed, N.E.; Vlková, M.; Faisal, M.B.; Silander, O.K. Rapid and Inexpensive Whole-Genome Sequencing of SARS-CoV-2 Using 1200 Bp Tiled Amplicons and Oxford Nanopore Rapid Barcoding. Biology Methods and Protocols 2021, 5, doi:10.1093/biomethods/bpaa014.

22. Rockett, R.J.; Arnott, A.; Lam, C.; Sadsad, R.; Timms, V.; Gray, K.-A.; Eden, J.-S.; Chang, S.; Gall, M.; Draper, J.; et al. Revealing COVID-19 Transmission in Australia by SARS-CoV-2 Genome Sequencing and Agent-Based Modeling. Nature Medicine 2020, 26, 1398–1404, doi:10.1038/s41591-020-1000-7.

23. Bolger, A.M.; Lohse, M.; Usadel, B. Trimmomatic: A Flexible Trimmer for Illumina Sequence Data. Bioinformatics 2014, 30, 2114–2120, doi:10.1093/bioinformatics/btu170.

24. Li, H.; Durbin, R. Fast and Accurate Short Read Alignment with Burrows-Wheeler Transform. Bioinformatics 2009, 25, 1754–1760, doi:10.1093/bioinformatics/btp324.

25. Grubaugh, N.D.; Gangavarapu, K.; Quick, J.; Matteson, N.L.; De Jesus, J.G.; Main, B.J.; Tan, A.L.; Paul, L.M.; Brackney, D.E.; Grewal, S.; et al. An Amplicon-Based Sequencing Framework for Accurately Measuring Intrahost Virus Diversity Using PrimalSeq and IVar. Genome Biology 2019, 20, 8, doi:10.1186/s13059-018-1618-7.

26. Turakhia, Y.; de Maio, N.; Thornlow, B.; Gozashti, L.; Lanfear, R.; Walker, C.R.; Hinrichs, A.S.; Fernandes, J.D.; Borges, R.; Slodkowicz, G.; et al. Stability of SARS-CoV-2 Phylogenies. PLOS Genetics 2020, 16, e1009175, doi:10.1371/JOURNAL.PGEN.1009175.

27. O’Toole, Á.; Scher, E.; Underwood, A.; Jackson, B.; Hill, V.; McCrone, J.T.; Colquhoun, R.; Ruis, C.; Abu-Dahab, K.; Taylor, B.; et al. Assignment of Epidemiological Lineages in an Emerging Pandemic Using the Pangolin Tool. Virus Evolution 2021, 7, doi:10.1093/VE/VEAB064.

28. Rambaut, A.; Holmes, E.C.; O’Toole, Á.; Hill, V.; McCrone, J.T.; Ruis, C.; du Plessis, L.; Pybus, O.G. A Dynamic Nomenclature Proposal for SARS-CoV-2 Lineages to Assist Genomic Epidemiology. Nature Microbiology 2020, 5, 1403–1407, doi:10.1038/S41564-020-0770-5.

29. Hueston, L.; Kok, J.; Guibone, A.; McDonald, D.; Hone, G.; Goodwin, J.; Carter, I.; Basile, K.; Sandaradura, I.; Maddocks, S.; et al. The Antibody Response to SARS-CoV-2 Infection. Open Forum Infectious Diseases 2020, 7, doi:10.1093/OFID/OFAA387.

30. SARS-CoV-2 Inactivation Testing: Interim Report; 2020;

31. Cele, S.; Jackson, L.; Khoury, D.S.; Khan, K.; Moyo-Gwete, T.; Tegally, H.; San, J.E.; Cromer, D.; Scheepers, C.; Amoako, D.G.; et al. Omicron Extensively but Incompletely Escapes Pfizer BNT162b2 Neutralization. Nature 2022, 602, 654–656, doi:10.1038/s41586-021-04387-1.

32. Arbel, R.; Hammerman, A.; Sergienko, R.; Friger, M.; Peretz, A.; Netzer, D.; Yaron, S. BNT162b2 Vaccine Booster and Mortality Due to Covid-19. N Engl J Med 2021, 1–8, doi:10.1056/NEJMoa2115624.

