## Supplementary material for "Improved neutralisation of the SARS-CoV-2 Omicron variant after Pfizer-BioNTech (BNT162b2) COVID-19 vaccine boosting with a third dose": Supplmentary Material

***Table S1: Study Participant Demographic and In-house SARS-CoV-2 Immunofluorescence (IFA) and Neutralising Antibody Titres (nAbT)***

| **Sera** | **Sex** | **Age**  **(yrs)** | **Timing post BNT162b2 (months)** | **SARS-CoV-2 IFA** | | | **Virus Lineage** | | |
| --- | --- | --- | --- | --- | --- | --- | --- | --- | --- |
|  |  |  |  | **IgG** | **IgA** | **IgM** | **Wildtype** | **Delta** | **Omicron** |
| **5** | M | 65 | 1 | 320 | <10 | <10 | 40 | 20 | <10 |
| **3** | M | 49 | 1 | 320 | <10 | <10 | 20 | <10 | <10 |
| **3** | M | 49 | 3 | 40 | <10 | <10 | <10 | <10 | <10 |
| **9** | F | 59 | 1 | 160 | <10 | <10 | <10 | <10 | <10 |
| **9** | F | 59 | 3 | 80 | <10 | <10 | <10 | <10 | <10 |
| **9** | F | 59 | Booster (1M) | 1280 | <10 | <10 | 320 | 320 | 160 |
| **1** | F | 36 | Naive | <10 | <10 | <10 | <10 | <10 | <10 |
| **1** | F | 36 | 3 | 80 | <10 | <10 | 40 | 20 | <10 |
| **14** | F | 34 | 6 | 80 | <10 | <10 | 20 | <10 | <10 |
| **14** | F | 34 | Booster (1M) | 640 | <10 | <10 | 160 | 160 | 40 |
| **15** | M | 63 | 6 | 40 | <10 | <10 | <10 | <10 | <10 |
| **15** | M | 63 | Booster (1M) | 640 | 20 | <10 | 160 | 80 | 80 |
| **16** | M | 59 | 6 | 40 | <10 | <10 | 20 | <10 | <10 |
| **16** | M | 59 | Booster (1M) | 160 | 20 | <10 | 320 | 160 | 40 |

**Key:** BNT162b2 – Pfizer-BioNTech (BNT162b2) vaccination; F – female; M – male; IFA – Immunofluorescence Assay; IgA – Immunoglobulin A; IgG – Immunoglobulin G; IgM – Immunoglobulin M; Omicron – Omicron (B.1.1.529) lineage; Wild-type – Wildtype (A.2.2) lineage; Delta – Delta (B.1.617.2) lineage; Booster (1M) – 4 weeks after 3rd booster dose of BNT162b2.

***Figure S1. Waning levels of SARS-CoV-2-specific IgG 1, 3 and 6 months following two doses of Pfizer-BioNTech (BNT162b2) which are boosted 4 weeks after the 3rd BNT162b2 dose (booster dose)***

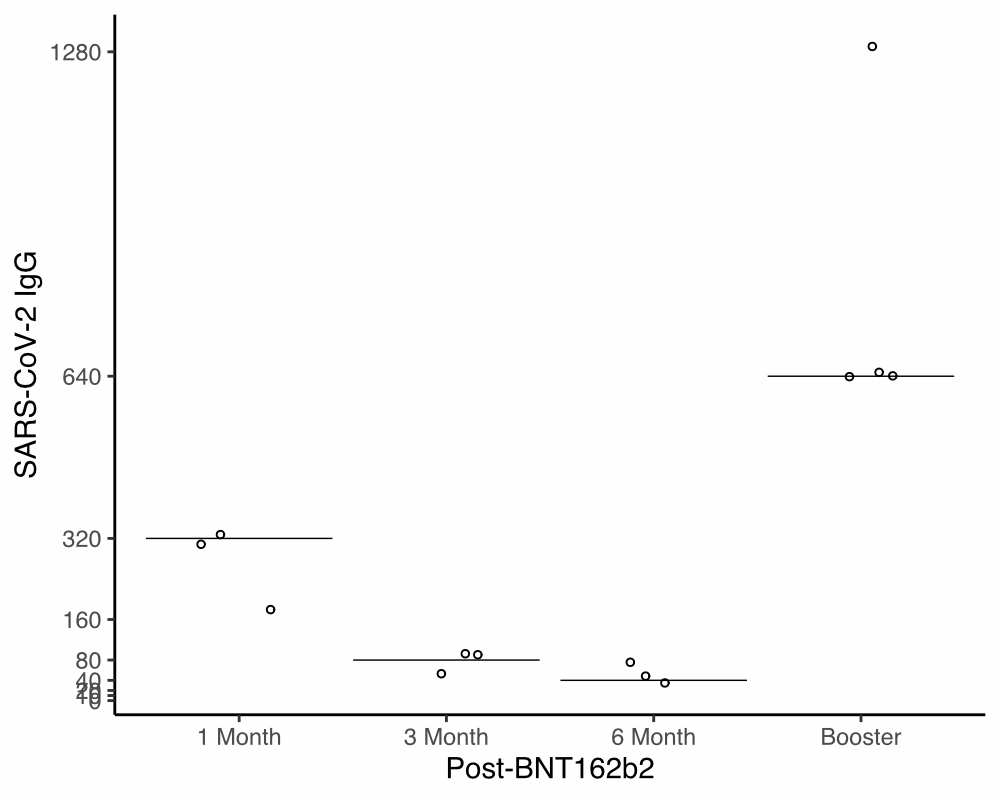

**Figure S3.** Illustrates levels of SARS-CoV-2-specific IgG 1, 3 and 6 months following two doses of Pfizer-BioNTech (BNT162b2) and 4 weeks after the 3rd dose (boosting dose) of BNT162b2. Levels have been determined using an immunofluorescence assay (IFA), black lines depict the median IgG level for each timepoint.

**Figure S2. *Infection kinetics of SARS-CoV-2 lineages propagated in VeroE6/TMPRSS2 cells***

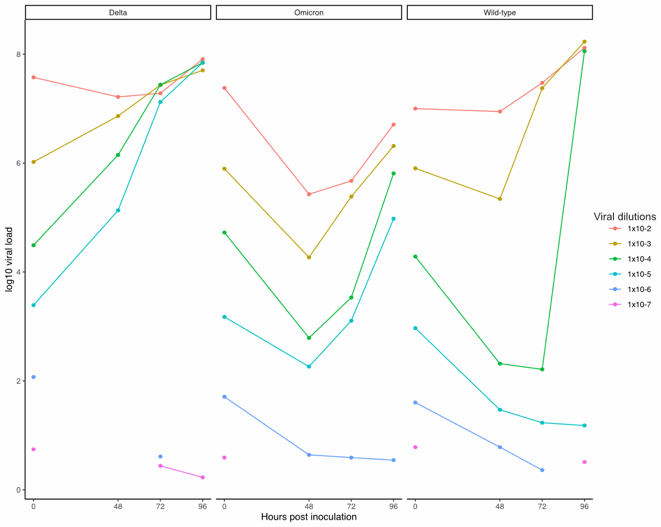

**Figure S2.** Illustrates infection kinetics of SARS-CoV-2 variants propagated in VeroE6/TMPRSS2 cells. Circles represent the viral load of serial dilutions of virus quantified at inoculation, 48, 72 and 96 hrs post-infection determined by SARS-CoV-2 in-house quantitative reverse transcriptase real time polymerase chain reaction (qRT-qPCR) targeting the *N*-gene

***Figure S3. Change in SARS-CoV-2 viral load 72 hours post-neutralisation with sera collected post- Pfizer-BioNTech (BNT152b2) challenged with SARS-CoV-2 VOCs Delta and Omicron compared to wildtype (lineage A.2.2) virus***

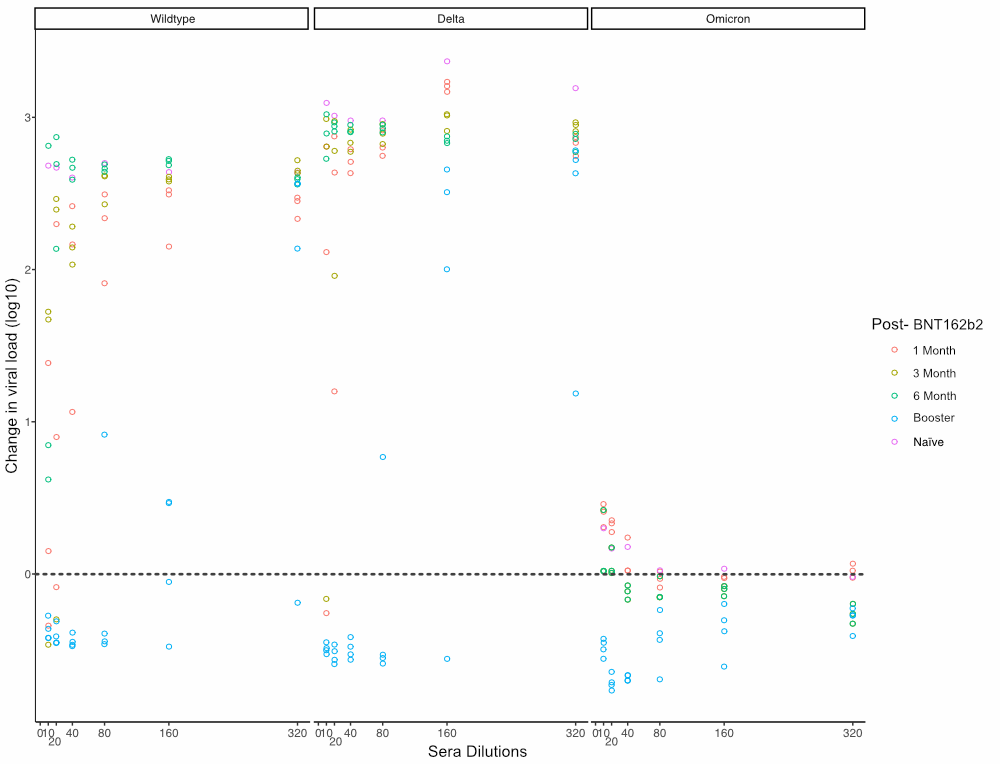

**Figure S3.** Illustrates replication of SARS-CoV-2 variants post-neutralisation when neutralised sera collected at different intervals post- Pfizer-BioNTech (BNT152b2). Dashed line indicates positive replication compared to the viral load determined for the viral inoculum by SARS-CoV-2 in-house quantitative reverse transcriptase real time polymerase chain reaction (qRT-qPCR) targeting the *N*-gene. Coloured circles represent sera collected 1 Month (red), 3 Months (yellow), 6 Months (green), 4 weeks after a booster dose (blue) and vaccine naïve sera (pink).

***Table S2: Non-Silent mutations in SARS-CoV-2 isolates used in Micro-neutralisation experiments.***

| **SARS-CoV-2 Virus** | **Non-silent Mutations** | **Pango Lineage**  **(GISAID accession)** |
| --- | --- | --- |
| **Omicron** | NSP3:K38R, NSP3:S1265(del), NSP3:A1892T, NSP4:T492I, NSP5:P132H, NSP6:L105(del), NSP6:I189V, NSP12b:P314L, NSP14:I42V, S:A67 (del), S:I68 (del), S:T95I, S:G142 (del), S:I210 (ins), S:N211(del), S:G339D, S:S371L, S:S373P, S:S375F, S:K417N, S:N440K, S:G446S, S:S477N, S:T478K, S:E484A, S:Q493R, S:G496S, S:Q498R, S:N501Y, S:Y505H, S:T547K, S:D614G, S:H655Y,S:N679K, S:P681H, S:N764K, S:D796Y*, S:N856K*, S:Q954H, S:N969K, S:L981F, E:T9I, M:D3G, M:Q19E, M:A63T, N:P13L*, N:E31(del)*, N:A134V*, N:RG203KR, ORF10:R24C | BA.1.17  (EPI_ISL_7987968) |
| **Delta** | NSP3:H323Y, NSP3:A488S, NSP3:P1228L, NSP3:P1469S, NSP4:V167L, NSP4:T492I, NSP6:T77A, NSP12b:P314L, NSP12b:G662S, NSP13:P77L, NSP14:A394V, NSP16:Q238H, S:T19R, S:T95I*, S:G142D*, S:E156(del), S:L452R, S:T478K, S:D614G, S:P681R, S:D950N,ORF3a:S26L,M:I82T, ORF7a:G70(del), ORF7a:V82A, ORF7a:T120I, ORF7b:T40I, ORF8:D119, N:D63G, N:R203M, N:G215C, N:D377Y | AY.39.1  (EPI_ISL_3398616) |
| **Wildtype** | NSP4:F308Y, ORF3a:G196V, ORF8:L84S, N:P13L, N:S197L | A.2.2  (EPI_ISL_427714) |

* Missing sequence data over mutations due to mismatches in primer sequences in one of the three SARS-CoV-2 lineages sequenced (Clinical specimen, 96 hrs post viral culture. 72 hrs post-neutralisation).
